# Analysis of Methicillin Resistance in *Staphylococcus Aureus* Sepsis Using TDbasedUFE

**DOI:** 10.1101/2024.01.25.577291

**Authors:** S. Watanabe, Y-h. Taguchi

## Abstract

scATAC-seq explains chromatin accessibility at cell-type resolution. Accordingly, this process is crucial for advancing our understanding of pathology and disease states. However, annotated data from scATAC-seq are both extensive and sparse; thus, conducting multidimensional analyses under multiple conditions is a challenging task. TDbasedUFE is a valuable tool for analyzing scATAC-seq data as it can extract genes in an unsupervised manner under multiple conditions based on tensor decomposition. We analyzed scATAC-seq data from the peripheral blood mononuclear cells of patients with sepsis infected with *S. aureus* using TDbasedUFE.

We extracted genes that exhibited different responses in methicillin-resistant (MSSA) and methicillin-sensitive *S. aureus* (MSSA) strains in sepsis for each cell type. Subsequently, we searched for studies containing gene sets similar to the extracted genes and predicted their functions. We also constructed protein-protein interactions (PPIs) for the extracted genes, defined hub proteins as central to the interactions based on degrees and clustering coefficients, and investigated the functions of these hub proteins. The genes of interest were abundant across all cell types, ranging from 710 to 1,372 genes. The functions of the extracted genes were predicted to be associated with several diseases or physiological substances. The hub proteins identified from the PPI analysis were mainly related to the ribosome, and their functions were associated with protein synthesis. These results highlight the suitability of TDbasedUFE for the analysis of scATAC-seq data. The functions of the genes identified in this study may provide insights into new promising therapeutic approaches, considering the distinction between methicillin resistance and *S. aureus* sepsis.

## Introduction

Gene expression is influenced by the chromatin structure. An Assay for Transposase Accessible Chromatin with high-throughput sequencing (ATAC-seq) provides a valuable profile for analyzing epigenetic regulatory functions by revealing the open chromatin regions^1^. Single-cell ATAC-seq (scATAC-seq) allows the examination of cellular heterogeneity by analyzing the cell type-specific distribution of chromatin accessibility^2^. However, as the total length of coverage of DNA fragments obtained from scATAC-seq is very short compared to the full length of the genome, the resulting scATAC-seq profile is very sparse. The scATAC-seq data under multiple conditions are formatted as high-dimensional tensors after sparse individual scATAC-seq profiles are aggregated, resulting in a difficult analysis. TDbasedUFE^3^ is a powerful application that applies tensor decomposition to analyze large tensors, enabling the extraction of biologically critical genes (e.g., disease-causing genes). Tensor decomposition has already been proven to be successful at analyzing large sparse matrices of scATAC-seq data^4^.

In a clinical setting, whether *Staphylococcus aureus* (*S. aureus*) is methicillin-resistant must be elucidated. In *S. aureus* infections, methicillin resistance serves as a guide for selecting an appropriate treatment option^5^, and the recommended antibiotics differ according to their methicillin-resistant *S. aureus* (MRSA) or methicillin-susceptible *S. aureus* (MSSA) status^6^. Similarly, in bacteremia caused by the same *S. aureus* infection as sepsis, early molecular diagnosis of MRSA is expected to reduce mortality and treatment costs^7^. However, the management of *S. aureus* in sepsis does not distinguish between MRSA and MSSA and antibiotics effective against MRSA are prophylactically recommended^8^. Furthermore, the increased frequency of vancomycin use as a first-line drug for MRSA infections is considered a contributing factor to the recent decrease in the vancomycin sensitivity of *S. aureus*^9^. Notably, empirical antibiotic therapy poses a public health risk. Therefore, comparing the epigenetic responses of patients with MRSA and MSSA infections in sepsis may lead to selective and more effective treatment strategies.

Distinguishing between MRSA and MSSA in sepsis has not been extensively achieved using the Differentially Expressed Genes (DEGs) obtained from the analysis of host single-cell RNA-sequencing (scRNA-seq) data^10^. The analysis of multi-omics data from scRNA-seq and scATAC-seq using MAGICAL revealed 53 circuit genes, which represents at most one-third of the circuit gene between infection and control^10^. The 53 circuit genes were expressed between those infected with MRSA and those infected with MSSA (area under the receiver operating characteristic curve from 0.67 to 0.75)^10^. In this study, using annotated scATAC-seq data^10^ obtained from peripheral blood mononuclear cell (PBMC) samples from healthy controls and patients infected with two strains of *S. aureus*, TDbasedUFE^3^ was employed to identify genes with distinct expression between MRSA and MSSA. Enrichment analysis was conducted using the obtained genes to determine their functions. Furthermore, using the STRING database^11^, we investigated the protein-protein networks to identify the hub proteins.

## Results

### Gene extraction by TDbasedUFE

We identified genes for the following six cell types: Naïve CD4^+^ T cells (CD4 Naive), CD4^+^ central memory T cells (CD4 TCM), CD8^+^ effector memory T cells (CD8 TEM), CD14^+^ monocytes (CD14 Mono), CD16^+^ monocytes (CD16 Mono), and natural killer cells (NK), using TDbasedUFE^3^. Table 1 lists the numbers of patients for each cell type. The condition was listed in Table 1. The patient counts correspond to the number of Singular Value Vectors (SVVs) in the bar graph under the same conditions (left red bar in Figure 1). Similarly, the right green bar graph in Figure 1 represents SVVs for each condition, where the conditions listed from left to right include the Control, MRSA, and MSSA. For all cell types, the index of condition *l*_3_ for SVV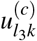with opposite signs for MRSA (*k* = 2) and MSSA (*k* = 3), was selected. In all cell types, SVVs under the same conditions (left red bar graph in Figure 1) had the same sign, indicating that tensor decomposition was performed to minimize the dependence on patients. In CD14 Mono, CD16 Mono, and CD4 Naive cells, the components of the control were the smallest, suggesting that genes distinguishing between MRSA and MSSA in healthy individuals were selected. CD4 TCM and NK cells had significantly larger absolute values specifically for MRSA, whereas in CD8 TEM, they were selected to differentiate between MSSA and the control. Although various independences of SSVs were found under the three conditions for individual cell types, all selected SSVs had large absolute values under the conditions of either MRSA or MSSA, indicating that the genes of interest were selected for all cell types (Figure 1).

**Table 1.** Sample for each cell type and infectious condition. The number of samples per cell type and per condition. The sample numbers were truncated to match the minimum value to ensure uniformity across conditions for analysis. Specific sample names and their attributes are summarized in Supplementary Table 1.

| CD14 Mono |  |  | CD16 Mono |  |  | CD4 Naive |  |  |
| --- | --- | --- | --- | --- | --- | --- | --- | --- |
| Control | MRSA | MSSA | Control | MRSA | MSSA | Control | MRSA | MSSA |
| 10 | 10 | 10 | 5 | 5 | 5 | 9 | 9 | 9 |

| CD4 TCM |  |  | CD8 TEM |  |  | NK |  |  |
| --- | --- | --- | --- | --- | --- | --- | --- | --- |
| Control | MRSA | MSSA | Control | MRSA | MSSA | Control | MRSA | MSSA |
| 10 | 10 | 10 | 10 | 10 | 10 | 5 | 5 | 5 |

**Figure 1.**
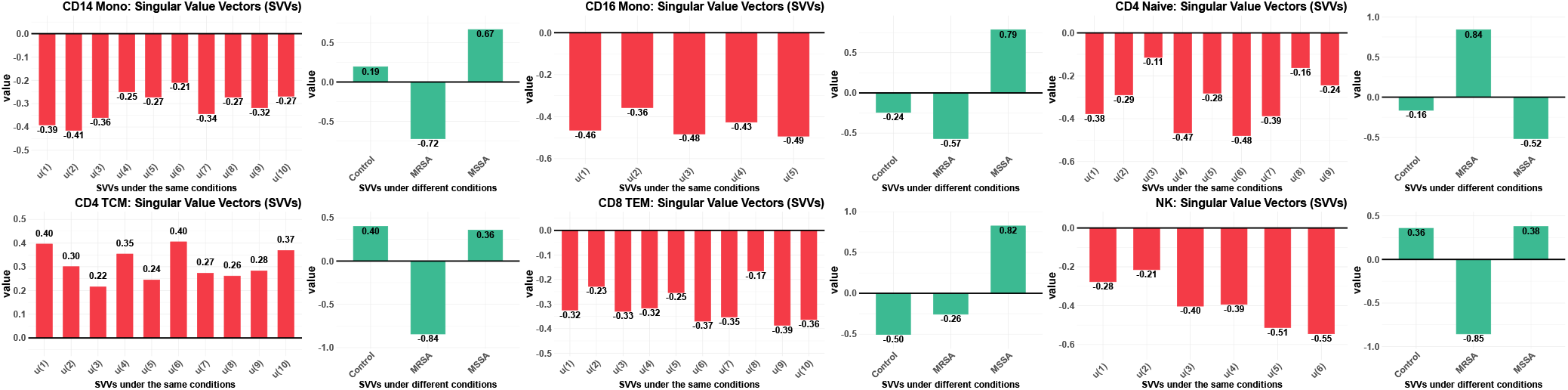
SVVs in the sample and condition for each cell type. The results of tensor decomposition for the six cell types (CD4 Naive, CD4 TCM, CD8 TEM, CD14 Mono, CD16 Mono, NK). The red bars on the left indicate the values of SVVs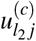in the sample direction; these values have the same sign in all cell types. The green bars on the right indicate the values of SVVs 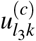 In the condition direction, with the largest difference between MRSA (*k* = 2) and MSSA (*k* = 3) selected. The SVVs were normalized, including their magnitude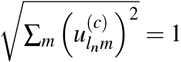. All SVVs in the sample direction and the condition directions are summarized in Supplementary Table 2.

The numbers of extracted genes and their similarities are summarized in Figure 2. Figure 2 (a) shows the number of genes commonly extracted from each cell type, where the diagonal elements represent the number of gene-specific cell types. The number of genes ranged from 710 for CD16 Mono to 1,372 for CD4 TCM. Figure 2 (b) shows the percentage of gene overlap between different cell types. These ratios are not symmetrical because they represent the proportion of genes present in the cell type of the column among those present in the cell type of the row. The largest gene overlap was observed between NK and CD8 TEM cells, with a similarity of 12%. This result suggests that the individual cell types share a limited number of genes with those selected for each cell type.

**Table 2.**
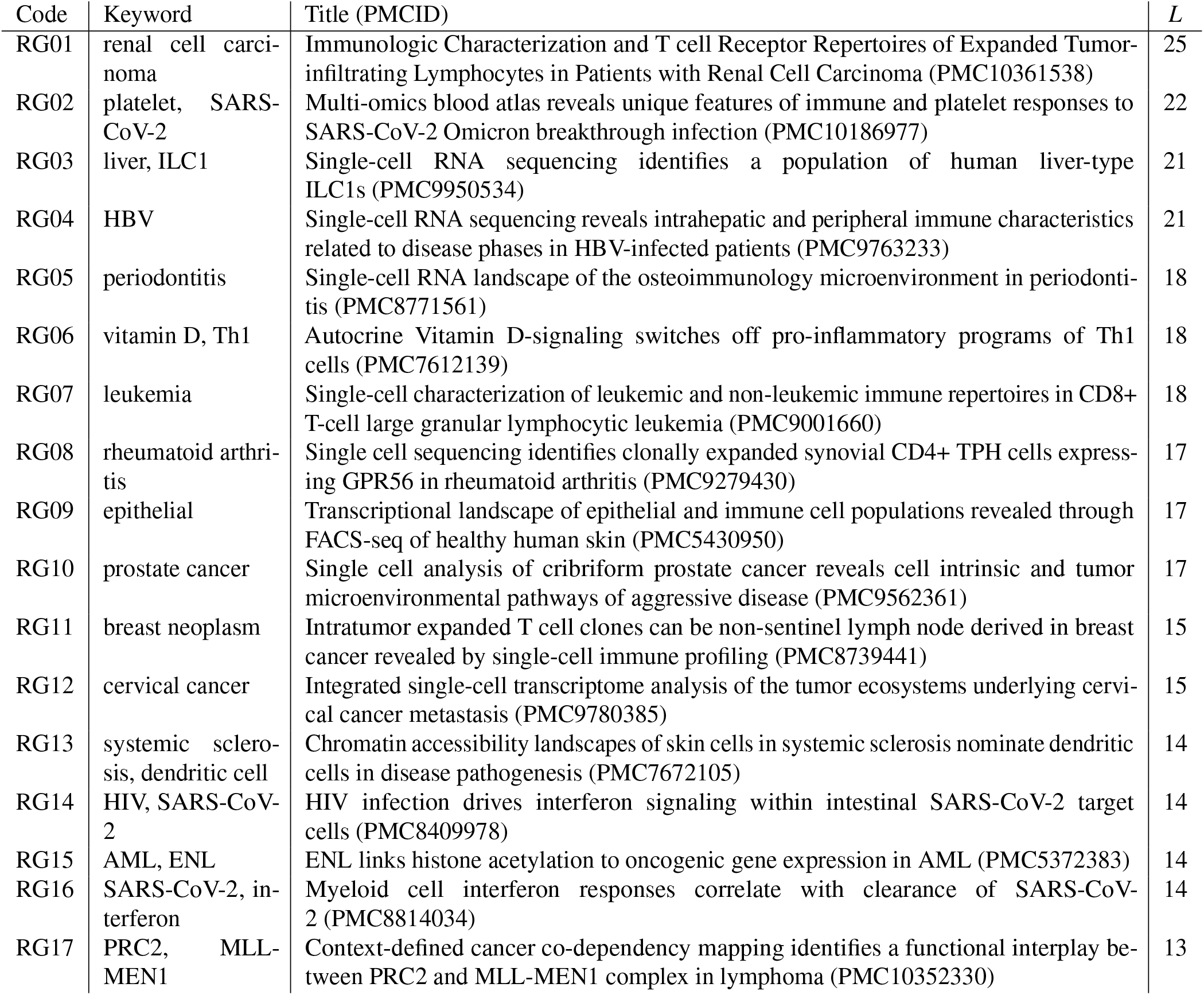
Paper and keywords for the top 1,000 selected gene sets across all cell types. Using Rummagene^12^, we extracted the top 1,000 terms with the smallest adjusted *P*-values for each cell type and selected across all cell types. *L* = ⟨log_10_ −*P*_ad_⟩ represents the average of the logarithm of adjusted *P*-values with reversed signs. A higher *L* value suggests a greater likelihood that the paper includes genes similar to our extracted genes. Keywords were extracted from the titles and abstracts of the papers, with a focus on terms related to disease names, organs, and other relevant aspects. Detailed results for the top 1,000 items in this table are available in Supplementary Table 3.

| Code | Keyword | Title (PMCID) | $L$ |
| --- | --- | --- | --- |
| RG01 | renal cell carcinoma | Immunologic Characterization and T cell Receptor Repertoires of Expanded Tumor-infiltrating Lymphocytes in Patients with Renal Cell Carcinoma (PMC10361538) | 25 |
| RG02 | platelet, SARS-CoV-2 | Multi-omics blood atlas reveals unique features of immune and platelet responses to SARS-CoV-2 Omicron breakthrough infection (PMC10186977) | 22 |
| RG03 | liver, ILC1 | Single-cell RNA sequencing identifies a population of human liver-type ILC1s (PMC9950534) | 21 |
| RG04 | HBV | Single-cell RNA sequencing reveals intrahepatic and peripheral immune characteristics related to disease phases in HBV-infected patients (PMC9763233) | 21 |
| RG05 | periodontitis | Single-cell RNA landscape of the osteoimmunology microenvironment in periodontitis (PMC8771561) | 18 |
| RG06 | vitamin D, Th1 | Autocrine Vitamin D-signaling switches off pro-inflammatory programs of Th1 cells (PMC7612139) | 18 |
| RG07 | leukemia | Single-cell characterization of leukemic and non-leukemic immune repertoires in CD8+ T-cell large granular lymphocytic leukemia (PMC9001660) | 18 |
| RG08 | rheumatoid arthritis | Single cell sequencing identifies clonally expanded synovial CD4+ TPH cells expressing GPR56 in rheumatoid arthritis (PMC9279430) | 17 |
| RG09 | epithelial | Transcriptional landscape of epithelial and immune cell populations revealed through FACS-seq of healthy human skin (PMC5430950) | 17 |
| RG10 | prostate cancer | Single cell analysis of cribriform prostate cancer reveals cell intrinsic and tumor microenvironmental pathways of aggressive disease (PMC9562361) | 17 |
| RG11 | breast neoplasm | Intratumor expanded T cell clones can be non-sentinel lymph node derived in breast cancer revealed by single-cell immune profiling (PMC8739441) | 15 |
| RG12 | cervical cancer | Integrated single-cell transcriptome analysis of the tumor ecosystems underlying cervical cancer metastasis (PMC9780385) | 15 |
| RG13 | systemic sclerosis, dendritic cell | Chromatin accessibility landscapes of skin cells in systemic sclerosis nominate dendritic cells in disease pathogenesis (PMC7672105) | 14 |
| RG14 | HIV, SARS-CoV-2 | HIV infection drives interferon signaling within intestinal SARS-CoV-2 target cells (PMC8409978) | 14 |
| RG15 | AML, ENL | ENL links histone acetylation to oncogenic gene expression in AML (PMC5372383) | 14 |
| RG16 | SARS-CoV-2, interferon | Myeloid cell interferon responses correlate with clearance of SARS-CoV-2 (PMC8814034) | 14 |
| RG17 | PRC2, MLL-MEN1 | Context-defined cancer co-dependency mapping identifies a functional interplay between PRC2 and MLL-MEN1 complex in lymphoma (PMC10352330) | 13 |

**Figure 2.**
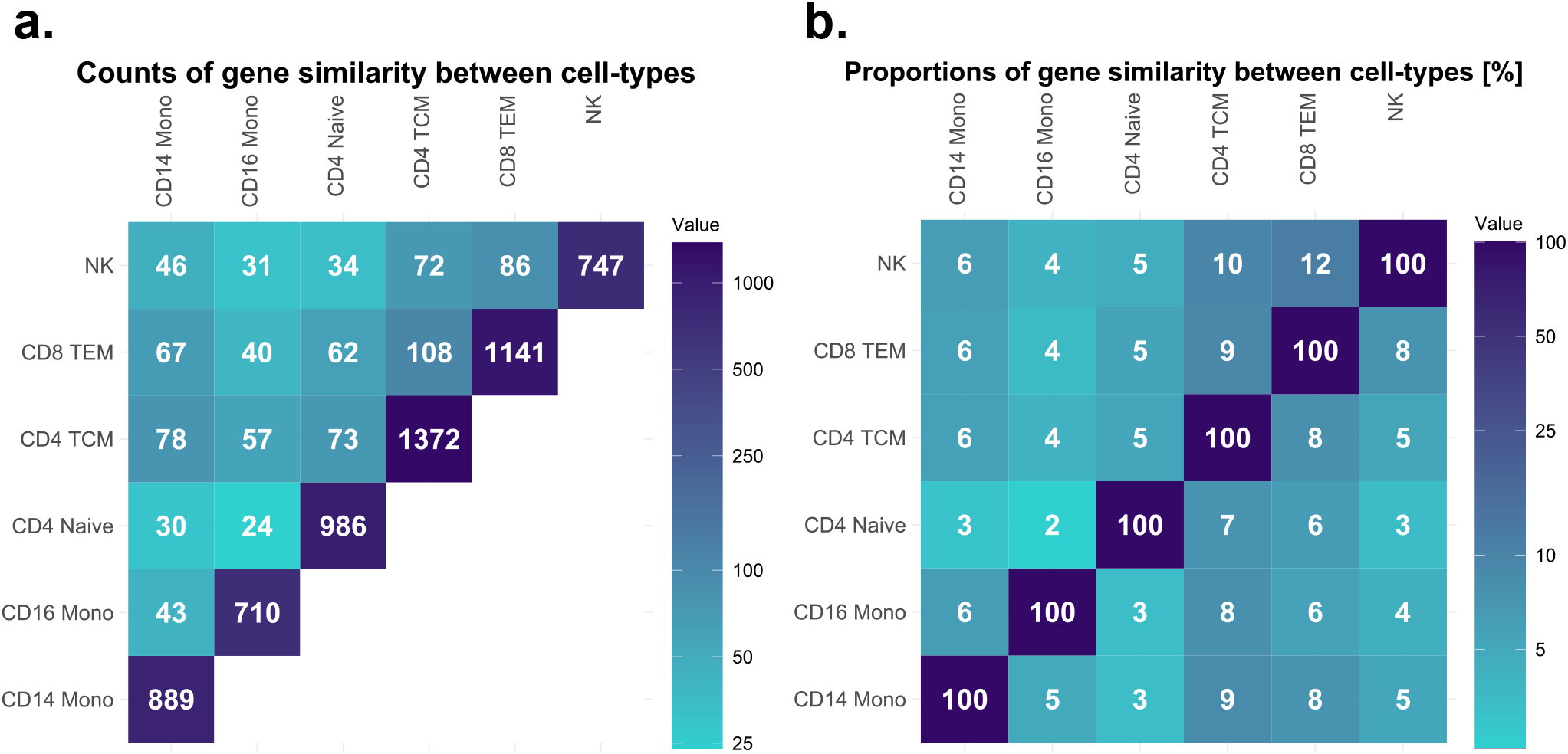
Counts and proportions of extracted genes and their similarity. **(a)** The counts of extracted genes and their similarity. This number represents the counts of genes selected together for each cell type. In particular, the elements on the diagonal represent the number of genes identified for that cell type. **(b)** The proportions of extracted genes and their similarity. This number represents the percentage of genes selected for the cell type on the horizontal axis that are also included in the cell type on the vertical axis (e.g. the “6” in the top-left corner indicates that 6% of the genes identified in NK cells are also identified in CD14 Mono cells). All extracted genes and SVVs in the gene direction are summarized in Supplementary Table 2.

### Enrichment analysis

In Table 2, papers that encompass gene sets selected across all cell types using Rummagene^12^, including their respective keywords, are listed. The criterion for selecting terms was based on the top 1,000 terms with smaller adjusted *P*-values for each cell type and common across all cell types. Thus, the papers in Table 2 were notably selected across all cell types, at least for the cell types in this study. The keywords were subjectively selected based on their biological importance from the titles or abstracts of each paper, with a maximum of two per paper.

The terms were assigned codes in increasing order of the geometric mean of *P*-values (or in decreasing order of *L*, which is the absolute value of logarithm of the adjusted *P*-value), with a higher *L* indicating a paper that is more likely to contain gene sets significantly resembling our extracted genes. These keywords predicted potential functions of the extracted genes related to the presence or absence of methicillin resistance in *S. aureus*.

The adjusted *P*-values for the top 1,000 terms ranged up to the 10^−7^ scale. The papers or gene sets listed in Table 2 had large *L* ≥ 13, indicating significant selection across all cell types. Among the gene sets involved in the study shown in Table 2, several gene sets with cell-type distinction were obtained from T cells and NK cells. Among the sequenced gene profiling methods, RNA-seq is the most frequently used. Keywords, such as tissue, disease, and molecular names, were found to markedly vary. Creating biologically meaningful classifications has become challenging owing to this diversity. Two keywords were commonly selected: “SARS-CoV-2” in RG02, RG14 and RG16, and “leukemia” in RG07 and RG15 (considering acute myeloid leukaemia (AML) as a type of leukemia). The detailed results for the top 1,000 items in this table are available in Supplementary Table 3.

The existence of papers supporting the association between keywords and methicillin resistance was summarized for each keyword in the subsequent “Discussion” section.

### STRING analysis

The extracted genes for each cell type were uploaded to STRING^11^ to obtain PPI data for each cell type. Only edges with a confidence score of > 0.900 were retained, capturing a network of strongly correlated proteins (Supplementary Figure 1). The average cluster coefficients^14^ ⟨*C*_*i*_⟩, which characterize the shape of the network and the total number of nodes representing the size of the network, are listed in Table 3. ⟨*C*_*i*_⟩ is the mean of all node cluster coefficients *C*_*i*_, with values close to 1 indicating a graph closer to a complete graph and values close to 0 indicating a graph closer to a tree graph^14^. ⟨*C*_*i*_⟩ for all cell types, except NK, falls within the range of 0.26±0.02, whereas NK exhibits an extremely low result of 0.14. Therefore, the PPI of NK had a structure similar to that of a tree, indicating relatively sparse and fewer interactions among proteins. The total number of nodes varied among different cell types, ranging from 155 for NK cells to 474 for CD4 TCM, indicating a wide range of node counts. Among the cell types, except for NK, with a similar ⟨*C*_*i*_⟩, the largest network had more than twice the number of nodes compared with the smallest network. NK had the smallest total number of nodes; however, the network size was not significantly correlated with ⟨*C*_*i*_⟩. Accordingly, NK cells may possess a specific PPI with a relatively small scale and sparse interactions. This result suggests that the functions of the gene products of the extracted genes in NK cells are relatively straightforward.

**Table 3.** Average cluster coefficient of PPI and the count of hub proteins for each cell type. The number of nodes, average cluster coefficient ⟨*C*_*i*_⟩, and the count of each hub protein for each cell type are listed. Hub proteins are proteins with a degree greater than five, which includes both normal hubs and family hubs. Normal hubs and family hubs are distinguished by their cluster coefficient ⟨*C*_*i*_⟩, with a ⟨*C*_*i*_⟩ *>* 0.5 being categorized as family hubs; otherwise, normal hubs. The “Rate” indicates the percentage of hub proteins that correspond to family hubs. The degree and clustering coefficient of all proteins are available in Supplementary Table 4. In addition, hub proteins can be classified based on any threshold selected using Supplementary Table 4.

| cell-type | Node | $\langle C_i \rangle$ | Hub Protein | Normal Hub | Family Hub | Rate [%] |
| --- | --- | --- | --- | --- | --- | --- |
| CD14 Mono | 223 | 0.25 | 8 | 8 | 0 | 00.0 |
| CD16 Mono | 222 | 0.25 | 9 | 7 | 2 | 22.2 |
| CD4 Naive | 256 | 0.25 | 11 | 10 | 1 | 09.1 |
| CD4 TCM | 474 | 0.28 | 71 | 46 | 25 | 35.2 |
| CD8 TEM | 334 | 0.26 | 33 | 22 | 11 | 33.3 |
| NK | 155 | 0.14 | 12 | 8 | 4 | 33.3 |

Table 3 lists the number of hub proteins for each cell. The hub type is presented in Table 3. Hub proteins were defined in the PPI network as proteins with a degree greater than five, indicating that they are central proteins interacting with a relatively large number of proteins. Among the hub proteins, those with a clustering coefficient greater than 0.5 (⟨*C*_*i*_⟩ *>* 0.5) were defined as “family hubs,” and others were categorized as “normal hubs” with ⟨*C*_*i*_⟩ ≤ 0.5 (Figure 3). Based on the definition of family hubs, these proteins form dense interaction groups, including those adjacent to each other. Thus, family hubs are the central proteins in a protein cluster connected by strong relationships.

**Figure 3.**
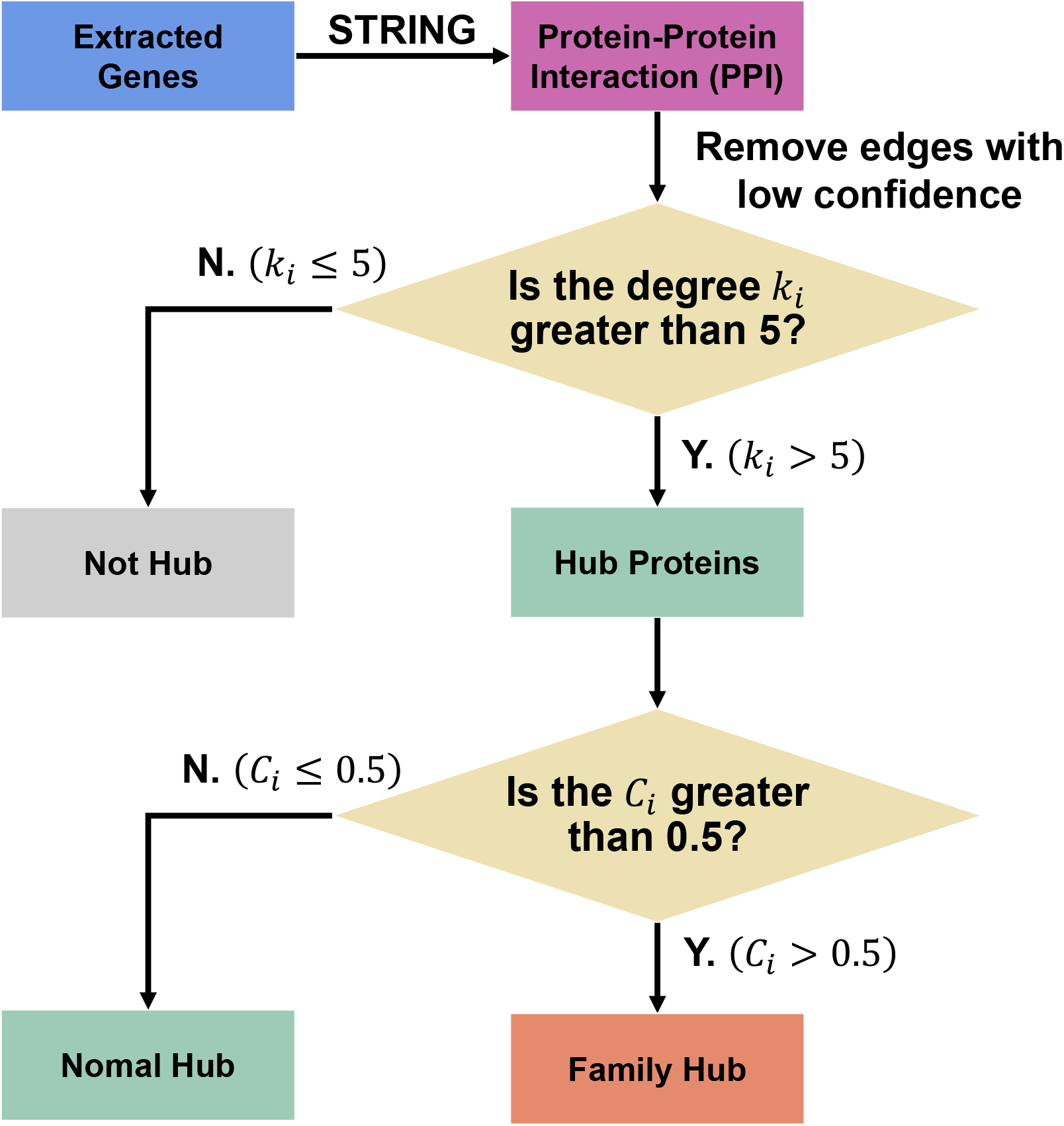
Flowchart from the generation of PPIs to the classification of hub proteins. PPIs were generated for each cell type using STRING^11^ with extracted genes using TDbasedUFE^3^. The PPI was filtered to retain only edges with a confidence greater than 0.900. Subsequently, isolated nodes were removed from the PPI. Proteins with a degree *k*_*i*_ greater than five were defined as hub proteins. Among these hub proteins, those with a cluster coefficient *C*_*i*_ *>* 0.5 were designated as family hubs, while those with a *C*_*i*_ ≤ 0.5 were classified as normal hubs.

The hub proteins in Table 3 indicate the total number of normal and family hubs, and the family rate represents the percentage of family hubs among the hub proteins. The number of nodes and counts of various hubs revealed a strong correlation, with a coefficient greater than 0.9. In contrast, the family rate did not show significant correlations with other values (with absolute value of correlation coefficients less than 0.7) and varied from 0% to 35% across the different cell types. Thus, the density of family hubs (or protein families) varies across cell types depending on the genes obtained for each cell type. In addition, despite the specifically low ⟨*C*_*i*_⟩ in NK, the family rate was 33% in the higher group. Therefore, no correlation was found between the family rate and network structure.

Table 4 lists the family hubs for each cell type, including their identifiers in approved symbols with HGNC nomenclature^13^. For CD8 TEM and CD4 TCM, a relatively large number of proteins was selected. However, a reduced number of unique identifiers was found due to the presence of several proteins associated with mitochondrial ribosomes (MRPL and MRPS) and ribosomal proteins (RPL and RPS). In NK cells, the four hub families were exclusively composed of ribosomal-related proteins. Figure 4 shows the network plots of genes and enriched terms for each library:`GO_Biological_Process_2023` (GOBP)^16^`GO_Cellular_Component_2023`, (GOCC)^16^ and, (KEGG)`KEGG_2021_Human`^17^. Combining all cell types, 16 terms were obtained for GOBP, 15 were obtained for GOCC, and 3 were obtained for KEGG. `GO_Molecular_Function_2023`(GOMF)^16^ terms were not obtained for any cell type.

**Table 4.** The family hubs obtained for each cell type. This table lists the proteins corresponding to the family hubs for each cell type, including their identifiers. Protein names are represented by approved symbols with HGNC nomenclature^13^, and identifiers are listed for proteins that appeared at least once. All family hubs are available in Supplementary Table 4.

| Cell-type | Protein | Identifier |
| --- | --- | --- |
| CD14 Mono |  |  |
| CD16 Mono | RPL14, RPL38 | RPL |
| CD4 Naive | MRPS10 | MRPS |
| CD4 TCM | ACTR5, EEFSEC, FBXW11, GTF2B, IMP3, MALSU1, MED21, MED9, MRPL1, MRPL16, MRPL39, MRPL41, MRPS14, MRPS18C, MRPS21, MRPS35, PSMA4, PSMB7, PSMB8, RPL10A, RPL18A, RPL26, RPL7L1, RPS27L, SRP54 | ACTR, EEFSEC, FBXW, GTF, IMP, MALSU, MED, MRPL, MRPS, PSMA, PSMB, RPL, RPS, SRP |
| CD8 TEM | EEF2, MRPL10, MRPL12, MRPL2, MRPL34, MRPL4, MRPL40, MRPL48, MRPL55, MRPS10, RPL26 | EEF, MRPL, RPL |
| NK | RPL23, RPS18, RPS27, RPS9 | RPL, RPS |

**Figure 4.**
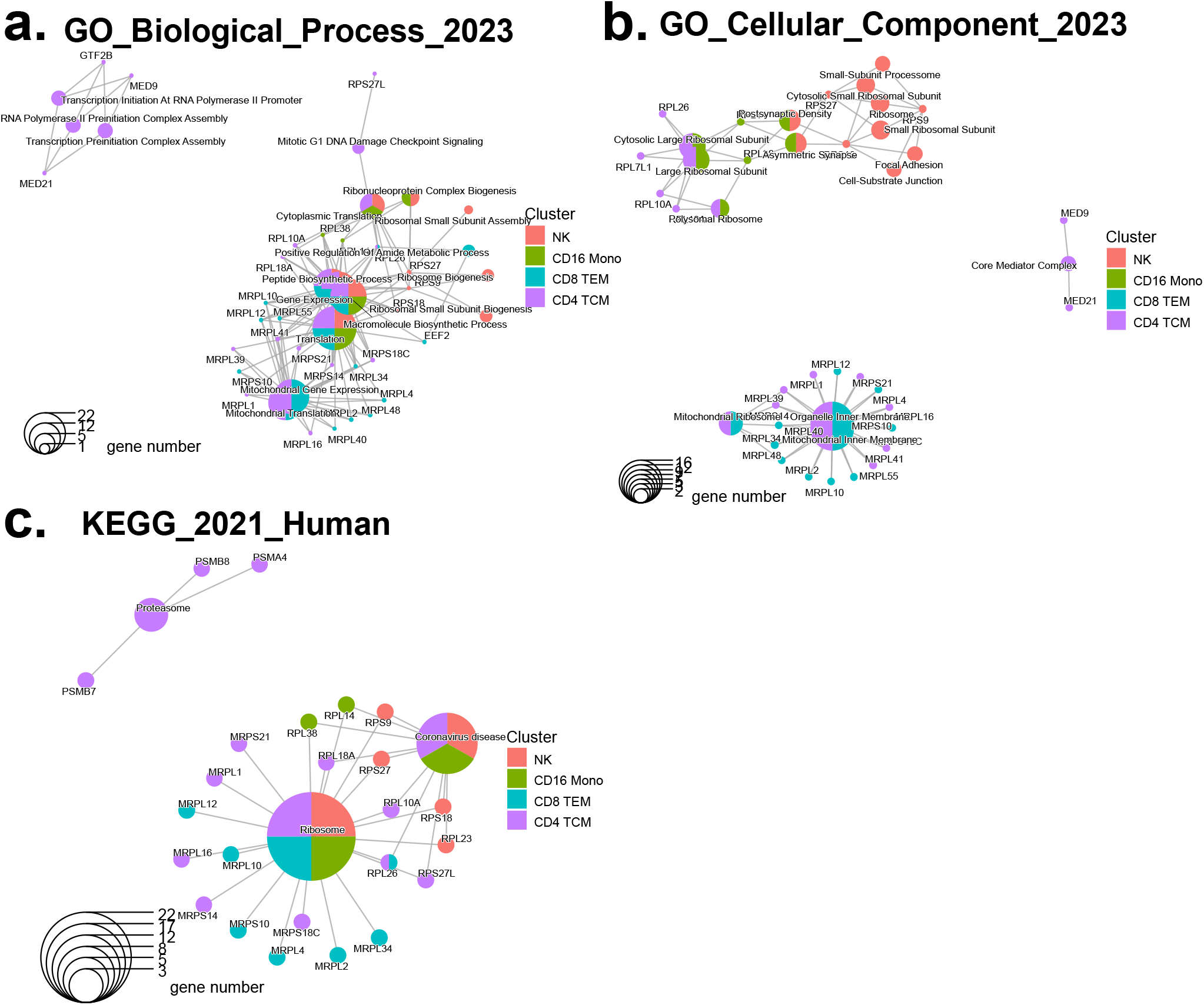
Results of the enrichment analysis for family hubs. The network was displayed using the `cnetplot()` function in clusterProfiler^15^. The network was constructed for each library: `GO_Biological_Process_2023` (GOBP)^16^ (a), `GO_Cellular_Component_2023` (GOCC)^16^ (b), and, `KEGG_2021_Human` (KEGG)^17^ (c). The selected enriched terms have adjusted *P*-values below the significance level of *α* = 0.01. The node colors correspond to cell types, and the mapping between colors and cell types is consistent across each network. The size of each term reflects the number of genes across all cell types. No `GO_Molecular_Function_2023` (GOMF)^16^ terms were obtained for any cell type. All results from the enrichment analysis of family hubs are available in Supplementary Table 5.

Multiple overlapping terms (nodes containing multiple colors) were observed across different cell types, suggesting that the biological roles of the family hub are shared among various cells. The abundance of terms related to gene expression, translation, and ribosomes GOBP suggests that the family hub plays a role in preparing for active protein synthesis. In GOCC, no terms were selected across three or more cell types, suggesting that the cellular activity of the family hub may vary among different cell types. In all libraries, terms associated with CD8 TEM were included in the terms for CD4 TCM, which are both memory T cell types. In GOCC, the terms from CD8 TEM formed a distinct network. Terms related to ribosomes were obtained for all cell types, providing further evidence that this family hub contributes to protein synthesis in all cell types. In KEGG, “Ribosomes” were obtained with very small *P*-values (ranging from *P*≈10^−12^ to *P*≈10^−7^). “Coronavirus disease” was the other term obtained for multiple cell types. All results of the enrichment analysis of the family hubs are available in the Supplementary Table 5.

## Discussion

This study sought to investigate the epigenetic functions of genes related to *S. aureus* sepsis by selecting genes from ATAC-seq data. The extracted genes represented the difference between the genetic responses of patients with sepsis infected with MRSA and those infected with MSSA, with the functions of these genes being unknown. The functions of the genes identified in this study may provide a novel perspective on the symptomatic treatment of *S. aureus* sepsis, considering differences in methicillin resistance. Enrichment analysis suggested several potential functions of the extracted genes and STRING analysis indicated potential functions related to ribosomes.

In gene extraction, owing to differences among patients for each cell type and the consistent direction of the SVVs under the same conditions, as indicated in Table 1, no patient bias was found in the identified genes. The genes were extracted based on genes that were significantly expressed (had high scores) in patients with MRSA and those with MSSA. Therefore, these genes are no longer genes uniquely expressed on one side (MRSA or MSSA).

The extracted genes were selected using scATAC-seq alone and a scoring metric for genes targeted at the Transcription Start Site (TSS)^10,18^. Therefore, gene ontologies and pathways may not function effectively as libraries for enrichment analysis due to this method. In the enrichment analysis, when subjecting the genes to Enrichr^19^ for each cell type, no significant terms were obtained from the GO library (GOBP, GOCC, GOMF) for all cell types. In the KEGG analysis, terms were only identified for memory T cells (CD4 TCM and CD8 TEM).

The relationship between the keywords in Table 2 obtained from enrichment analysis and methicillin resistance in *S. aureus* is supported by several papers found during the analysis (Table 5). The keywords identified in these previous studies strongly suggest the functions of the extracted genes, providing evidence that the approach used in this study with Rummage^12^ successfully predicted the gene functions. Keywords can serve as clinical markers for determining methicillin resistance in *S. aureus* that causes sepsis or as clues when exploring diseases, in which methicillin resistance in *S. aureus* could be a risk factor.

**Table 5.** Claims supporting the association between keywords and the MRSA versus MSSA comparison. Papers claiming an association between the keywords mentioned in Table 2 and the MRSA versus MSSA comparison are listed in this table. The code is consistent with that in Table 2. Keywords were only listed if relevant papers were found.

| Code | Keyword | Claim of the Paper |
| --- | --- | --- |
| RG02 | platelet | MRSA bacterial growth was significantly higher in platelet-rich plasma when compared to MSSA (*in vitro*)<sup>20</sup>. |
| RG04 | HBV | A statistically significant positive correlation between HCV (+) and MSSA carrier ($P = 0.001$) was observed<sup>21</sup>. |
| RG08 | rheumatoid arthritis | MRSA septic arthritis has a tropism for the glenohumeral joint while MSSA has a tropism for the knee<sup>22</sup>. |
| RG16 | interferon | MRSA strain can induce an exacerbated immune response in retinal cells (*in vitro*)<sup>23</sup>. |

## Methods

An overview of these approaches is presented in Figure 5.

**Figure 5.**
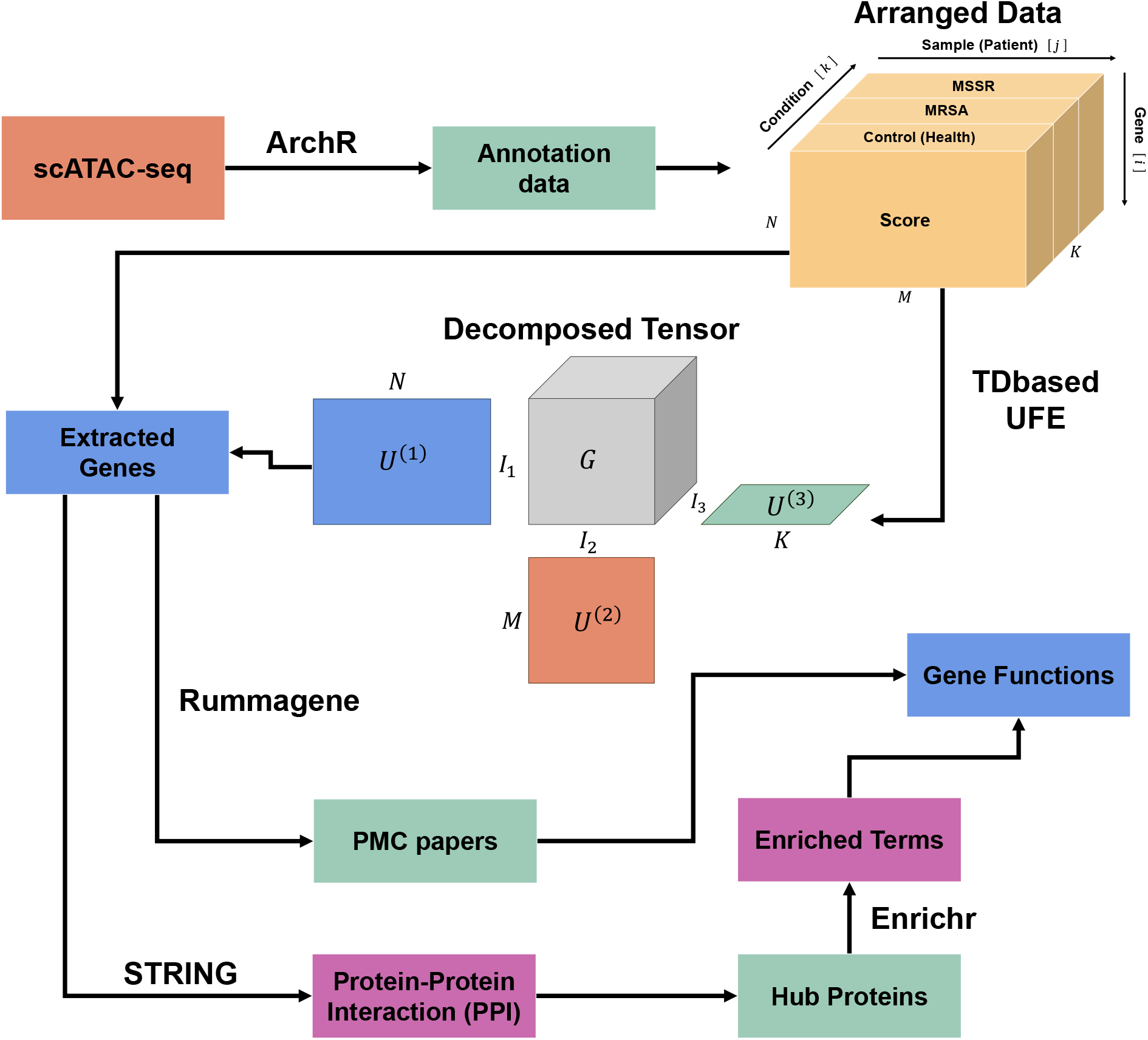
Brief Overview of the Entire Process. The analysis was conducted using the available `GSE220188` dataset on Gene Expression Omnibus (GSO).`GSE220188` includes scATAC-seq data from PBMC annotated with ArchR^18^ for patients with sepsis and healthy controls. The annotation data were categorized by cell type, and the fragment scores were arranged into third-order tensors aligned with three axes genes, samples, and patient conditions. The tensors created for each cell type were subjected to tensor decomposition using TDbasedUFE^3^. From the resulting vectors, genes exhibiting significant differences between MRSA and MSSA were extracted. The extracted genes were searched in PMC papers using Rummagene^12^ to identify papers containing similar gene sets. Papers ranked within the top 1,000 for each cell type and chosen across all cell types were selected. Subsequently, we investigated the relationship between keywords in PMC papers and methicillin resistance. We also predicted the functions of the genes. In addition, using STRING analysis^11^, hub proteins expected to play crucial roles in important functions were identified. We classified hub proteins into two categories based on their clustering coefficient: highly interconnected family hubs and normal hubs with weaker interaction connections. We conducted enrichment analysis using Enrichr^19^ with the Gene Ontology (GO)^17^ and KEGG pathway (KEGG)^24^ libraries for genes with the same name as the family hub proteins.

### Data set

The dataset is included in series `GSE220188`^10^ of the Gene Expression Omnibus (GSO) repository provided by the National Center for Biotechnology Information (NCBI). `GSE220188`^10^ is scATAC-seq that has already been annotated using ArchR^18^ of PBMCs from 10 MRSA samples, 11 MSSA samples, and 23 healthy control samples. In this study, the analysis was conducted only on cell types derived from the MRSA, MSSA, and healthy control groups, ensuring a sample size of five or more for each category (the sample count for each condition and each cell type). Specific sample names and their attributes are summarized in Supplementary Table 1.

### Tensor decomposition

We opted to provide an explanation of TDbasedUFE^3^. Briefly, for each cell type, we arranged the scored datasets into third-order tensors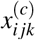 with three axes: gene, sample, and infectious conditions. Let *N* be the number of genes. In addition, for a specific cell type, let *M*_*c*_ denote the number of samples and *K*_*c*_ the number of infectious condition types. In this study, three conditions were considered: healthy controls, MRSA, and MSSA, resulting in *K*_*c*_ = 3. Furthermore, the number of genes was *N* = 23746. The number of patients *M*_*c*_ is provided in Table 1. Let 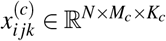 represent the score assigned to the *i*th gene of the *j*th sample under the *k*th condition for a certain cell type (its index in *c*). The tensors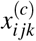were standardized using the gene vector 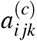from the scored scATAC-seq data for each *j* and *k*: Therefore, the average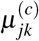and distribution 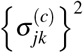 of 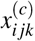 were

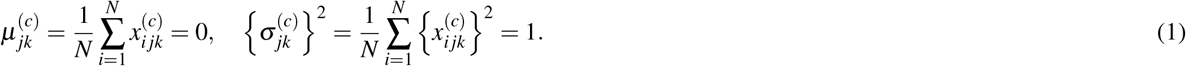

When applying higher-order singular value decomposition (HOSVD)^3,24^to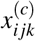using the gene size of the core tensor *I*_1_ ∈ ℕ, the following result is obtained

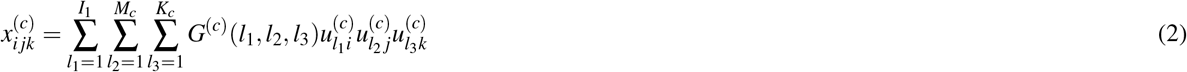

where 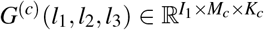 is a core tensor representing the weight of the product 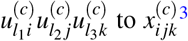. 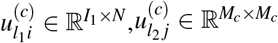, and 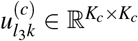are singular value orthogonal matrices (SVOM)^3^ (Figure 6 (a)). Choosing a value for *l*_*n*_, (*n* = 1, 2, 3) results in the selection of a vector from 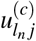, which is referred to as the Singular Value Vector (SVV) that represents differences in the corresponding direction^3^ (for example, when considering 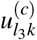 representing conditions, if a difference exists between 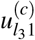and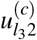, a difference between the first and second conditions is implied). *I*_1_ ∈ ℕ, (*I*_1_ ≤ *N*) also represents the multilinear rank^24^ in the gene direction (its index in *i*). To simplify the calculations, we set *I*_1_ = 10. The other ranks were not omitted for subsequent “Unsupervised feature extraction,” as they are square matrices (Figure 6 (a)).

**Figure 6.**
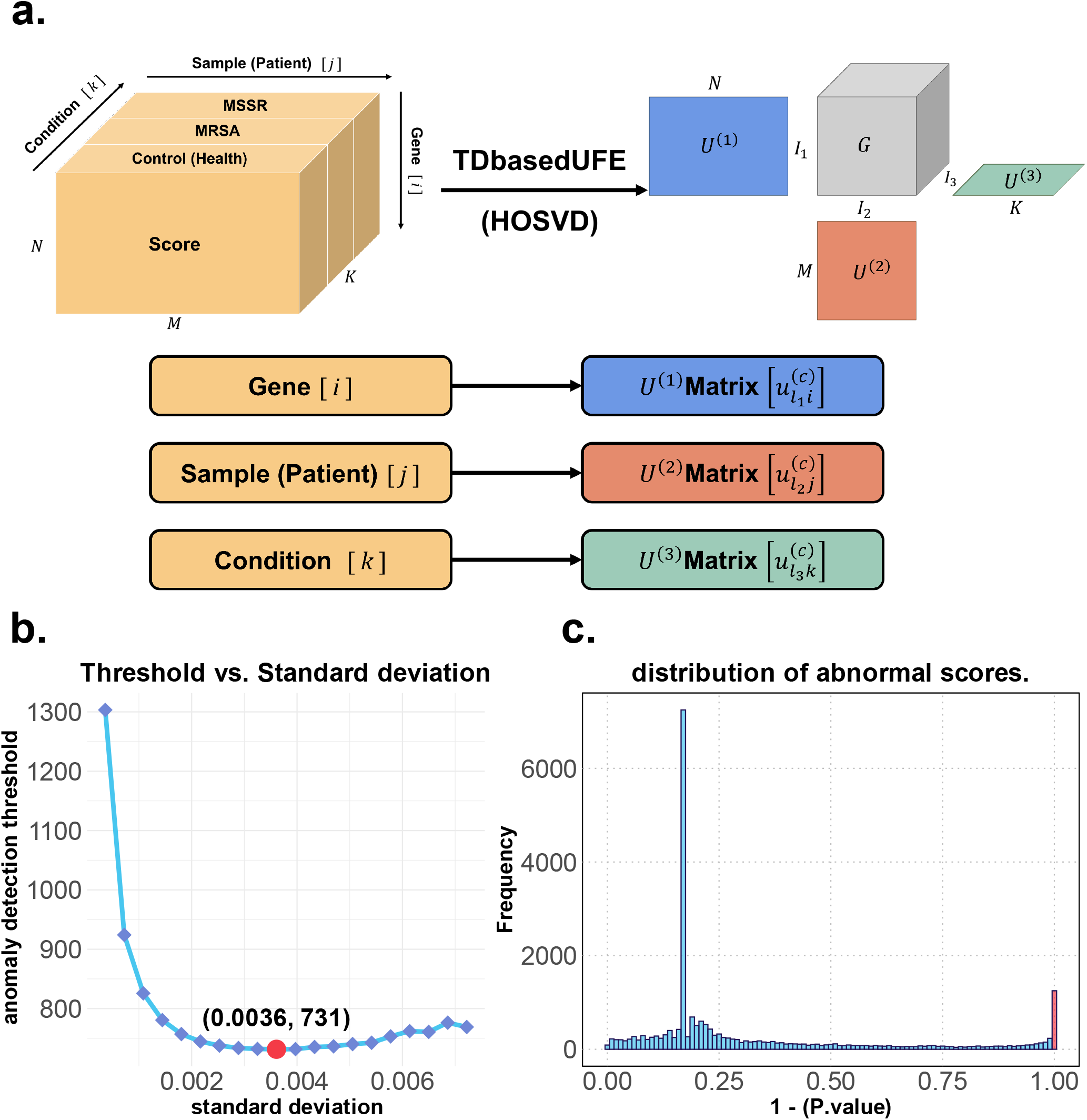
TDbasedUFE ‘s input and output. **(a)** TDbasedUFE^3^ used HOSVD^3,24^to decompose the score tensor into three SVOMs 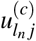, each with the same number of tensor dimensions. The resulting SVOMs correspond to the directions of the input tensor, such as Gene, Sample, or Condition. **(b)** This is an example of *c* = CD14 Mono. Standard deviation and the corresponding threshold. The x-axis indicates the standard deviation 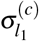 while the y-axis indicates the standard deviation of the histogram extracted through a *χ*^2^ test based on 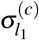. The red points represent the points where the threshold becomes the minimum value; this was adopted as the optimized standard deviation 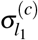 in equation (3)^25^. **(c)** This is also an example of *c* = CD14 Mono. In the histogram of 1 − *Q*_*i*_ with the optimized standard deviation 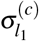, the red bar on the right represents the selected genes with *Q*_*i*_ *<* 0.01.

### Unsupervised feature extraction

The tensor decomposition of TDbasedUFE^3^ allows unsupervised feature extraction. However, a decomposition method must be chosen for our study. In this study, we created tensors with three conditions: control (*k* = 1), MRSA (*k* = 2), and MSSA (*k* = 3) to determine the differences between MRSA and MSSA. Accordingly, we chose *l*_3_, where the largest difference exists between 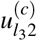 and 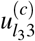. In contrast, to minimize the differences among patients, we chose *l*_2_ where all 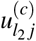values have the same sign. Under these conditions, *l*_2_ and *l*_3_ were uniquely determined in this analysis. Thereafter, *l*_1_ was selected to maximize the absolute value of core tensor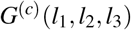. Consequently, we obtained the SVV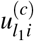, which represents the different genes identified under the condition of methicillin resistance of *S. aureus*. Assuming that the obtained SVV distribution followed a Gaussian distribution (null hypothesis), we selected the genes that exhibited significant differences. As a 2-tailed *χ*^2^ test was conducted, the *P*-value *P*_*i*_ is

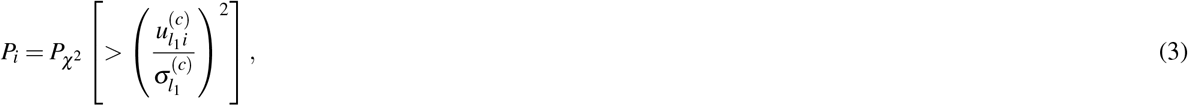

where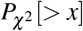 is the cumulative *χ*^2^ distribution and the argument is larger than *x*^3^. 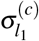 is the optimized standard deviation, such that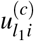obeys a Gaussian distribution as much as possible^25^ (Figure 6 (b)). Thereafter, the obtained *P*-values *P*_*i*_ were adjusted using the Benjamini-Hochberg (BH) procedure^26^. 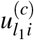with adjusted *P*-values *Q*_*i*_ below a threshold were selected, and these were considered as the extracted genes (Figure 6 (c)). In this study, the threshold was set to 0.01.

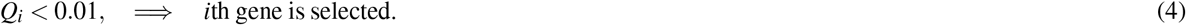

### Enrichment analysis

We proceeded to evaluate the gene sets that exhibited similarities to the extracted genes. The genes extracted from each cell type were processed using Rummagene^12^ and their overlap with publicly available gene sets in PubMed Central (PMC) was verified. In this gene set, as more PMC gene sets were selected using Rummagene^12^, we considered the terms chosen for all cell types to represent the functions associated with the extracted genes. The top 1,000 terms with the smallest *P*-values, adjusted using the BH procedure^26^, were selected for each cell type. Subsequently, we extracted the terms chosen for all cell types.

The relevance to genes was assessed by taking the score, *L*, which is the logarithm base 10 of the geometric mean of the adjusted *P*-values (or the average of the logarithm base 10 of the adjusted *P*-values *P*_ad_), represented by the following formula:

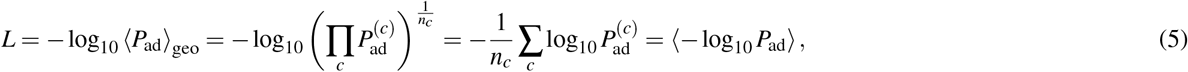

where *n*_*c*_ represents the total number of cell types, that is, *n*_*c*_ = 6, as shown in Table 1. In addition, ⟨*X* ⟩_geo_ denotes the geometric mean of *X*, and 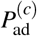denotes the adjusted *P*-value for the terms upon input of the genes extracted from the *c*th cell type.

Overall, papers with a larger *L* are more likely to include gene sets closely related to those extracted. Keywords, such as disease and tissue names, were extracted from the terms selected by Rummagene^12^, and their association with methicillin resistance was investigated.

### STRING analysis

We conducted STRING analysis^11^ for each cell type. Protein-protein interactions (PPI) were considered when confidence, representing the credibility of the interaction, was greater than 0.900. Nodes that were not connected to any other node (degree 0) were excluded from the PPI network. We analyzed the graph-theoretical properties of the obtained PPI using the R package “igraph”^27^ and identified hub proteins that could be central to the protein interactions. The definition of a hub is based on the study by Jing-Dong J. Han et al.^28^, in which a node with a degree greater than five is considered a hub protein. Maslov and Sneppen argued that proteins that bind to hub proteins do not tend to interact with other hub proteins^29^. Therefore, in this PPI, where only strong correlations were retained, the boundaries between the hub proteins were expected to become clearer. A cluster formed by interactions centered around a hub is referred to as a community. We calculated the clustering coefficient for each hub to measure the extent to which interactions occurred within communities. As proposed by Watts and Strogatz^14^, the clustering coefficient is defined as the proportion of actual connections among neighboring nodes. Therefore, the clustering coefficient *C*_*i*_ is computed as

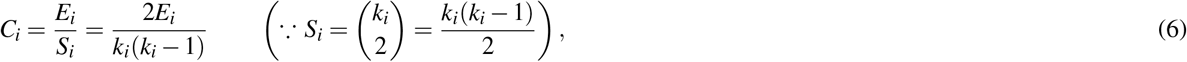

where *k*_*i*_ ≥2 is the *i*th node of degree. *E*_*i*_ is the number of connections among neighboring nodes, and *S*_*i*_ is the number of possible connections among neighboring nodes. Among hub proteins, those with a clustering coefficient *C*_*i*_ *>* 0.5 were defined as “family hubs.” In contrast, hub proteins with a cluster coefficient *C*_*i*_ ≤0.5 were defined as “normal hubs” (Figure 3).

These hub proteins are speculated to have communities that resemble complete graphs and are likely to possess biological functions. Furthermore, by definition, family hubs are likely to overlap with gene or protein families, indicating that they are part of a family in a PPI context. We identified hubs, particularly crucial family hubs, using the network obtained from the STRING analysis.

Subsequently, to investigate the biological functions of the family hub obtained, gene set enrichment analysis was performed using Enrichr^19^ for genes with the same names as those in the family hub. We conducted enrichment analysis for each cell type using four libraries of Enrichr^19^: `GO_Biological_Process_2023` (GOBP)^16^`GO_Cellular_Component_2023`, (GOCC)^16^, `GO_Molecular_Function_2023`(GOMF)^16^, `KEGG_2021_Human` and, (KEGG)^17^. The enrichment analysis was performed using the R package “clusterProfiler”^15^, with a significance level of *α* = 0.01. *P*-values were corrected using the BH procedure^26^.

## Supporting information

Supplementary Figure 1: All PPIs for each cell type.

## Acknowledgements

We would like to thank Editage for English language editing.

## Author contributions statement

Y.T. conceived the study, S.W. and Y.T. designed the research methods, and S.W. analyzed the data. All authors reviewed the manuscript.

## References

1. Buenrostro, J. D., Giresi, P. G., Zaba, L. C., Chang, H. Y. & Greenleaf, W. J. Transposition of native chromatin for fast and sensitive epigenomic profiling of open chromatin, DNA-binding proteins and nucleosome position. Nat. methods 10, 1213–1218, DOI: 10.1038/nmeth.2688 (2013).

2. Buenrostro, J. D. et al. Single-cell chromatin accessibility reveals principles of regulatory variation. Nature 523, 486–490, DOI: 10.1038/nature14590 (2015).

3. Taguchi, Y.-h. & Turki, T. TDbasedUFE and TDbasedUFEadv: bioconductor packages to perform tensor decomposition based unsupervised feature extraction. bioRxiv 2023–05, DOI: 10.1101/2023.05.14.540687 (2023).

4. Taguchi, Y.-h. & Turki, T. Tensor decomposition discriminates tissues using scATAC-seq. Biochimica et Biophys. Acta (BBA)-General Subj. 1867, 130360, DOI: 10.1016/j.bbagen.2023.130360 (2023).

5. Liu, C. et al. Clinical practice guidelines by the Infectious Diseases Society of America for the treatment of methicillin-resistant Staphylococcus aureus infections in adults and children. Clin. infectious diseases 52, e18–e55, DOI: 10.1093/cid/ciq146 (2011).

6. Kimmig, A. et al. Management of Staphylococcus aureus Bloodstream Infections. Front. Medicine 7, DOI: 10.3389/fmed.2020.616524 (2021).

7. Brown, J. & Paladino, J. A. Impact of rapid methicillin-resistant Staphylococcus aureus polymerase chain reaction testing on mortality and cost effectiveness in hospitalized patients with bacteraemia: a decision model. Pharmacoeconomics 28, 567–575, DOI: 10.2165/11533020-000000000-00000 (2010).

8. Evans, L. et al. Surviving Sepsis Campaign: International Guidelines for Management of Sepsis and Septic Shock 2021. Critical Care Medicine 49, e1063–e1143, DOI: 10.1097/ccm.0000000000005337 (2021).

9. Hiramatsu, K. Vancomycin-resistant staphylococcus aureus: a new model of antibiotic resistance. The Lancet Infect. Dis. 1, 147–155, DOI: 10.1016/s1473-3099(01)00091-3 (2001).

10. Chen, X. et al. Mapping disease regulatory circuits at cell-type resolution from single-cell multiomics data. Nat. Comput. Sci. 3, 644–657, DOI: 10.1038/s43588-023-00476-5 (2023).

11. Szklarczyk, D. et al. The STRING database in 2021: customizable protein–protein networks, and functional characterization of user-uploaded gene/measurement sets. Nucleic acids research 49, D605–D612, DOI: 10.1093/nar/gkaa1074 (2021).

12. Clarke, D. J. et al. Rummagene: Mining Gene Sets from Supporting Materials of PMC Publications. bioRxiv 2023–10, DOI: 10.1101/2023.10.03.560783 (2023).

13. Gray, K. A., Seal, R. L., Tweedie, S., Wright, M. W. & Bruford, E. A. A review of the new HGNC gene family resource. Hum. genomics 10, 1–9, DOI: 10.1186/s40246-016-0062-6 (2016).

14. Watts, D. J. & Strogatz, S. H. Collective dynamics of ‘small-world’ networks. nature 393, 440–442, DOI: 10.1038/10.1038/30918 (1998).

15. Yu, G., Wang, L.-G., Han, Y. & He, Q.-Y. clusterProfiler: an R package for comparing biological themes among gene clusters. Omics: a journal integrative biology 16, 284–287, DOI: 10.1089/omi.2011.0118 (2012).

16. Gene Ontology, C. et al. The gene ontology knowledgebase in 2023. Genetics 224, iyad031, DOI: 10.1093/genetics/iyad031 (2023).

17. Kanehisa, M., Furumichi, M., Tanabe, M., Sato, Y. & Morishima, K. KEGG: new perspectives on genomes, pathways, diseases and drugs. Nucleic acids research 45, D353–D361, DOI: 10.1093/nar/gkw1092 (2017).

18. Granja, J. M. et al. ArchR is a scalable software package for integrative single-cell chromatin accessibility analysis. Nat. genetics 53, 403–411, DOI: 10.1038/s41588-021-00790-6 (2021).

19. Kuleshov, M. V. et al. Enrichr: a comprehensive gene set enrichment analysis web server 2016 update. Nucleic acids research 44, W90–W97, DOI: 10.1093/nar/gkw377 (2016).

20. López, C., Álvarez, M. E., Carmona, J. U. et al. Temporal bacteriostatic effect and growth factor loss in equine platelet components and plasma cultured with methicillin-sensitive and methicillin-resistant Staphylococcus aureus: a comparative in vitro study. Vet. medicine international 2014, DOI: 10.1155/2014/525826 (2014).

21. Aydogan, U. et al. To study the correlation between carrier status of nasal Staphylococcus aureus in patients on haemodialysis with hepatitis C, hepatitis B and their sociodemographic features. West indian medical journal 61 (2012).

22. Al-Nammari, S. S., Bobak, P. & Venkatesh, R. Methicillin resistant Staphylococcus aureus versus methicillin sensitive Staphylococcus aureus adult haematogenous septic arthritis. Arch. orthopaedic trauma surgery 127, 537–542, DOI: 10.1007/s00402-007-0285-z (2007).

23. Naik, P. & Joseph, J. Difference in Host Immune response to Methicillin-Resistant and Methicillin Sensitive Staphylococcus aureus (MRSA and MSSA) Endophthalmitis. Ocular Immunol. Inflamm. 30, 1044–1054, DOI: 10.1080/09273948.2020.1859551 (2022).

24. De Lathauwer, L., De Moor, B. & Vandewalle, J. A multilinear singular value decomposition. SIAM journal on Matrix Analysis Appl. 21, 1253–1278, DOI: 10.1137/S0895479896305696 (2000).

25. Taguchi, Y.-h. & Turki, T. Principal component analysis-and tensor decomposition-based unsupervised feature extraction to select more suitable differentially methylated cytosines: Optimization of standard deviation versus state-of-the-art methods. Genomics 115, 110577, DOI: 10.1016/j.ygeno.2023.110577 (2023).

26. Benjamini, Y. & Hochberg, Y. Controlling the false discovery rate: a practical and powerful approach to multiple testing. J. Royal statistical society: series B (Methodological) 57, 289–300, DOI: 10.1111/j.2517-6161.1995.tb02031.x (1995).

27. Csárdi, G. et al. igraph: Network Analysis and Visualization in R, DOI: 10.5281/zenodo.7682609 (2023). R package version 1.5.1.

28. Han, J.-D. J. et al. Evidence for dynamically organized modularity in the yeast protein–protein interaction network. Nature 430, 88–93, DOI: 10.1038/nature02555 (2004).

29. Maslov, S. & Sneppen, K. Specificity and stability in topology of protein networks. Science 296, 910–913, DOI: 10.1126/science.1065103 (2002).

