## Supplementary figures and images for "Analysis of Methicillin Resistance in *Staphylococcus Aureus* Sepsis Using TDbasedUFE"

### Supplementary_Figure_1.pdf

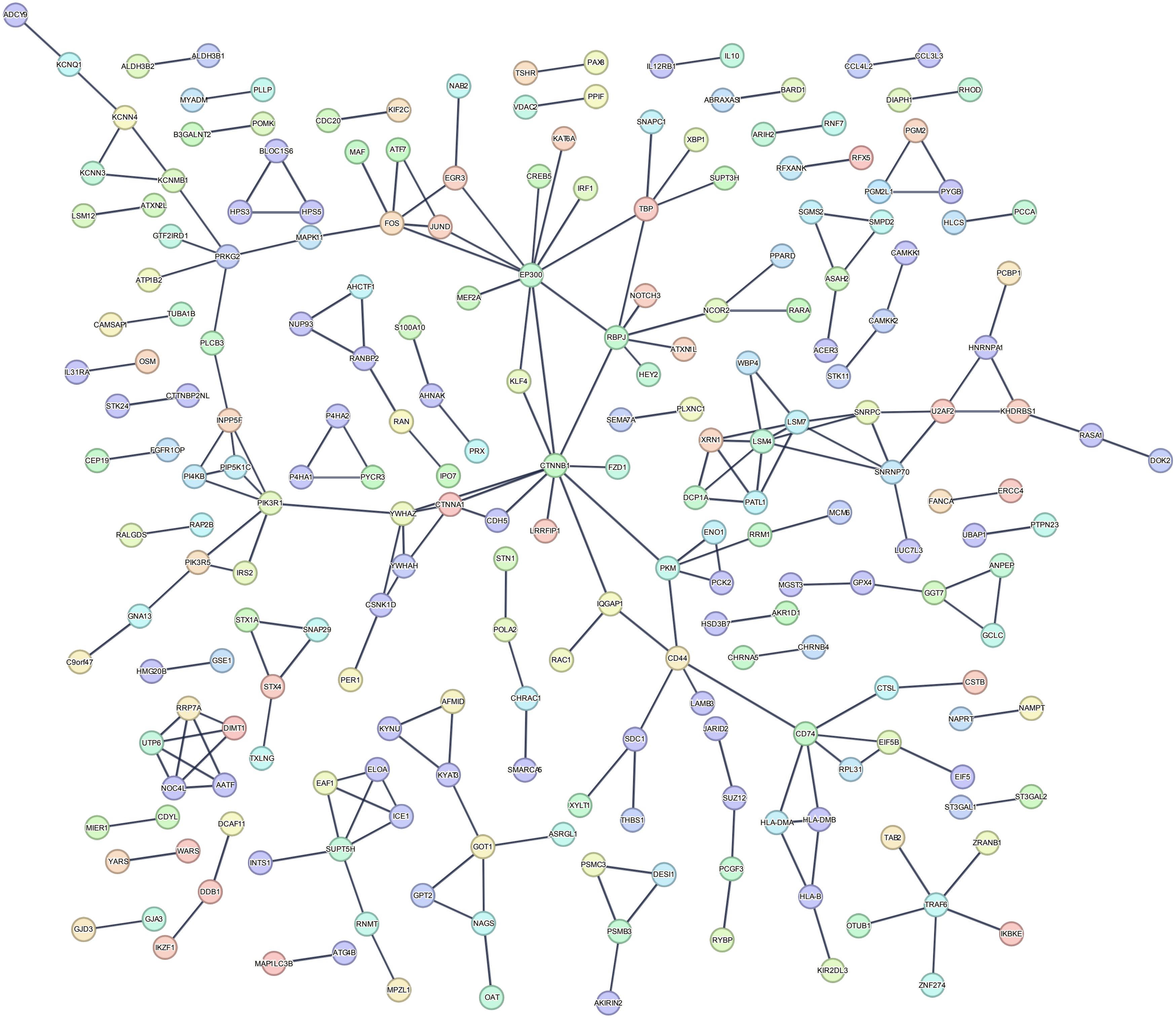

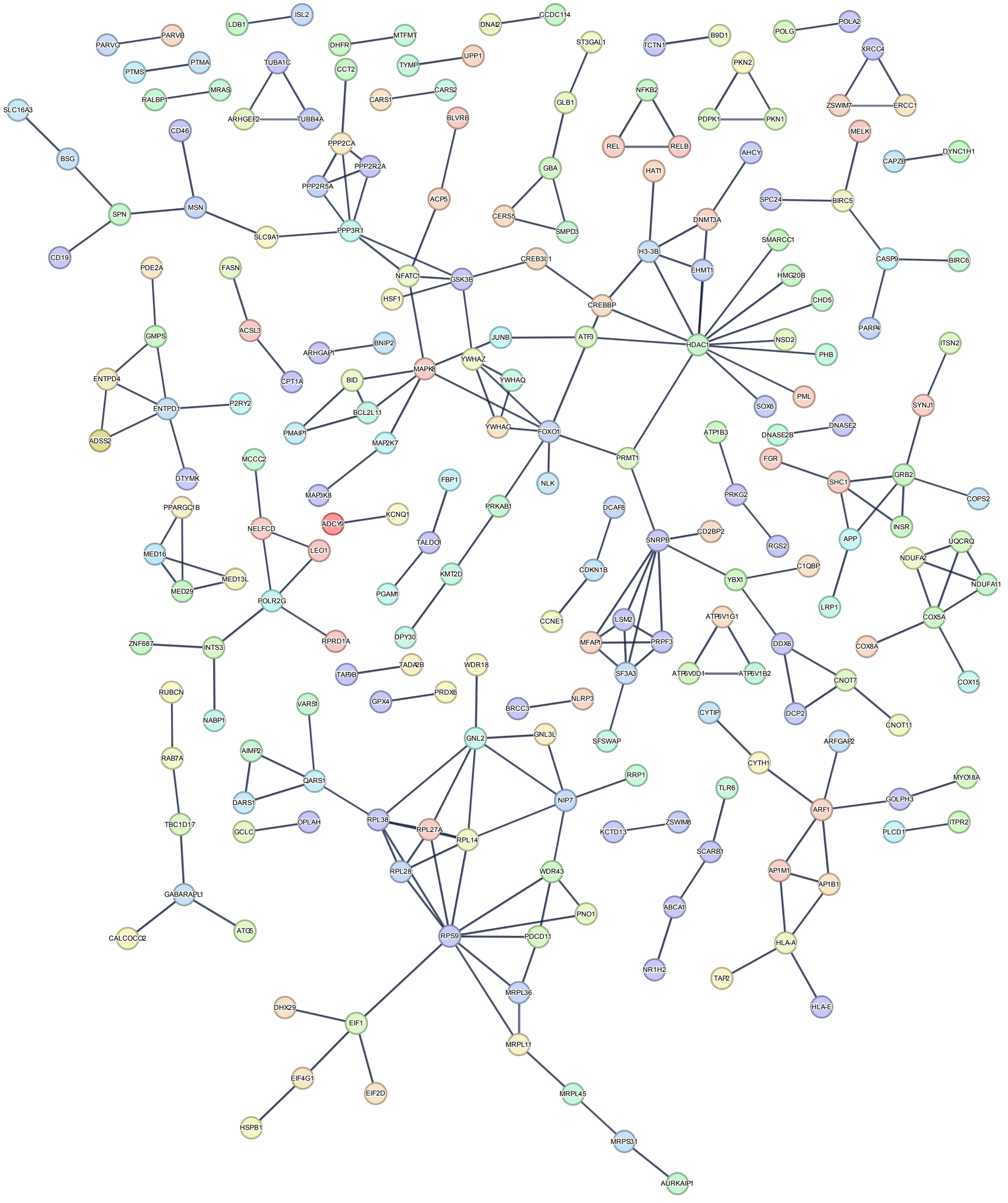

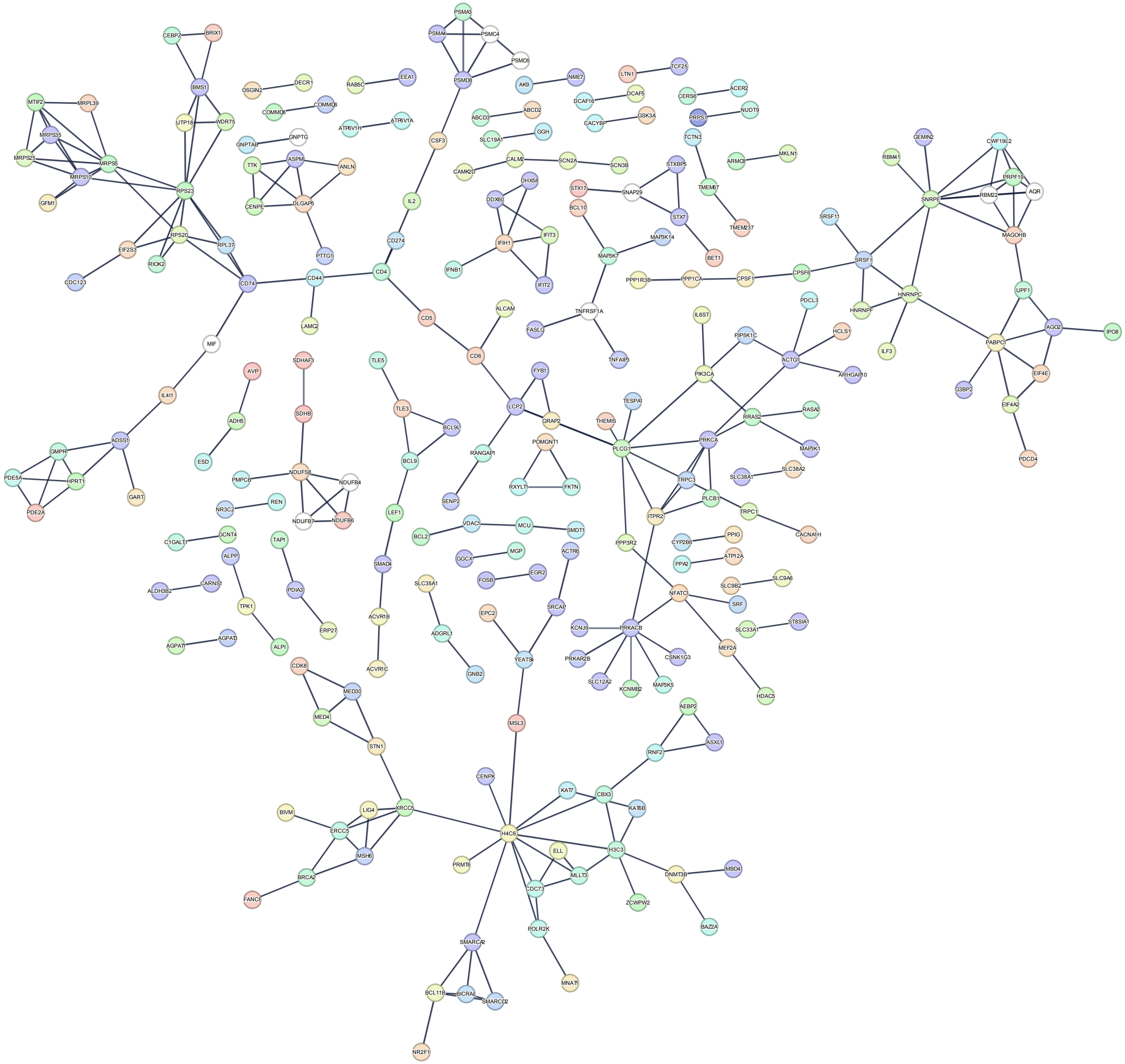

#### d. CD4 TCM

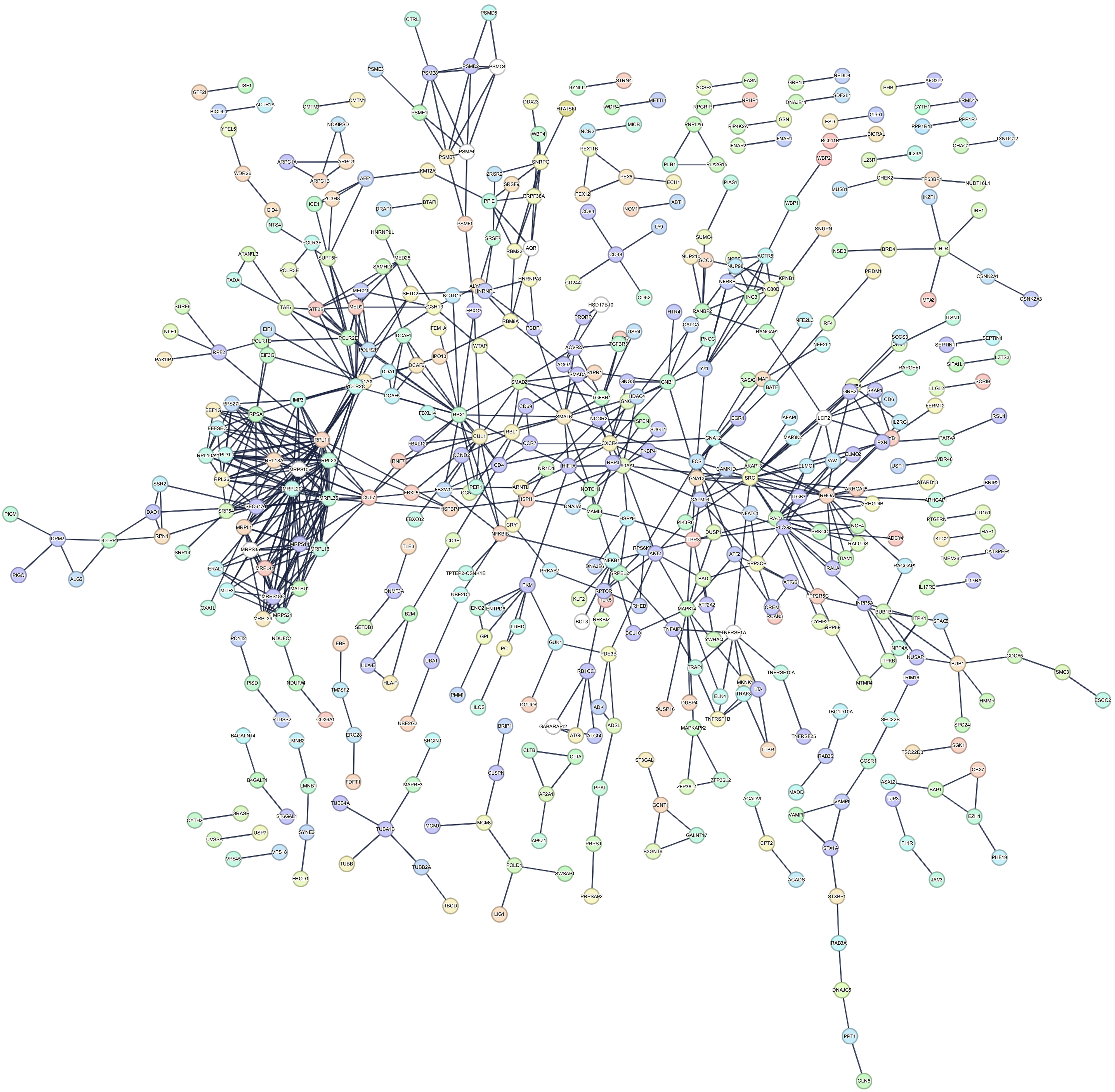

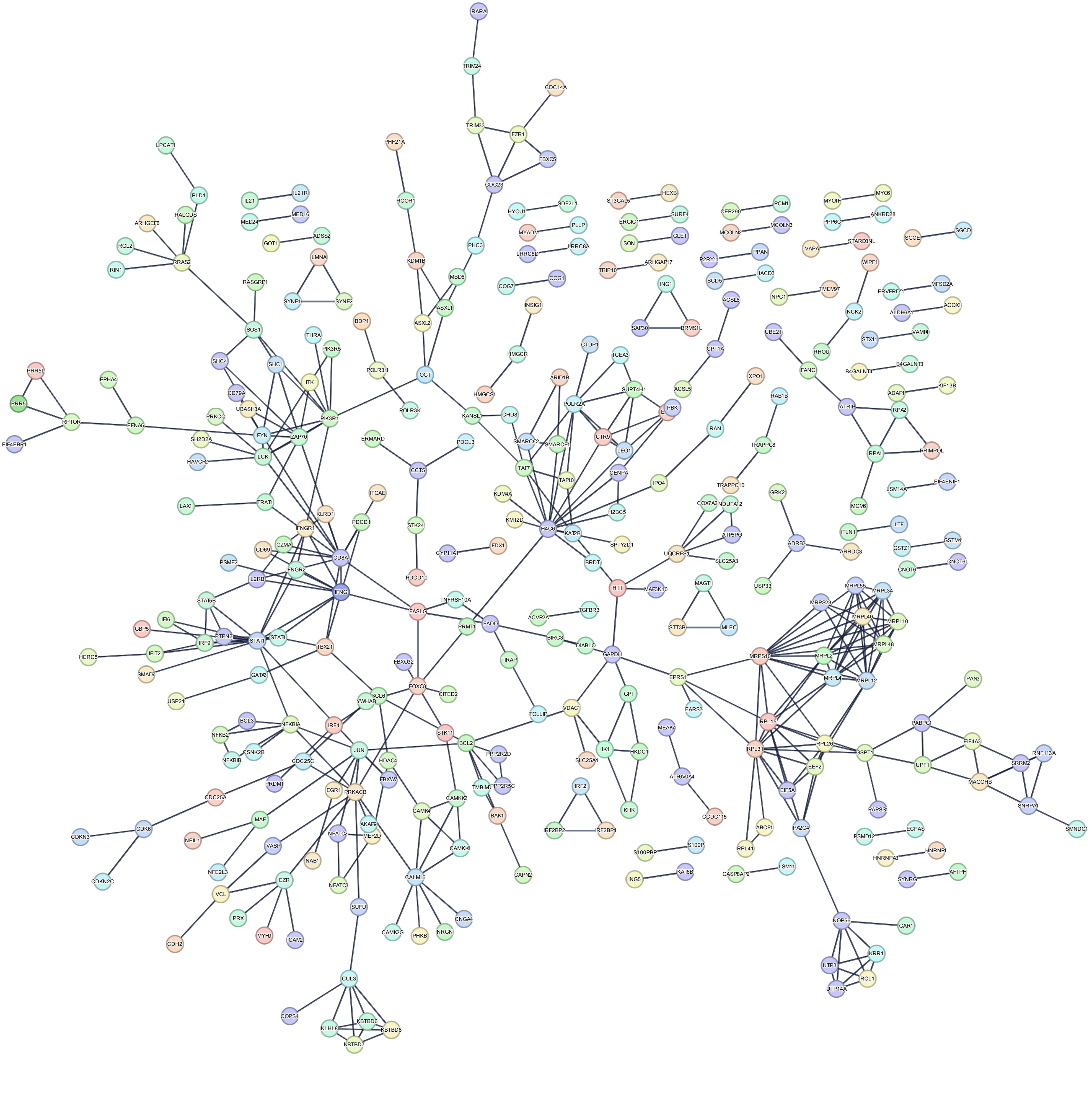

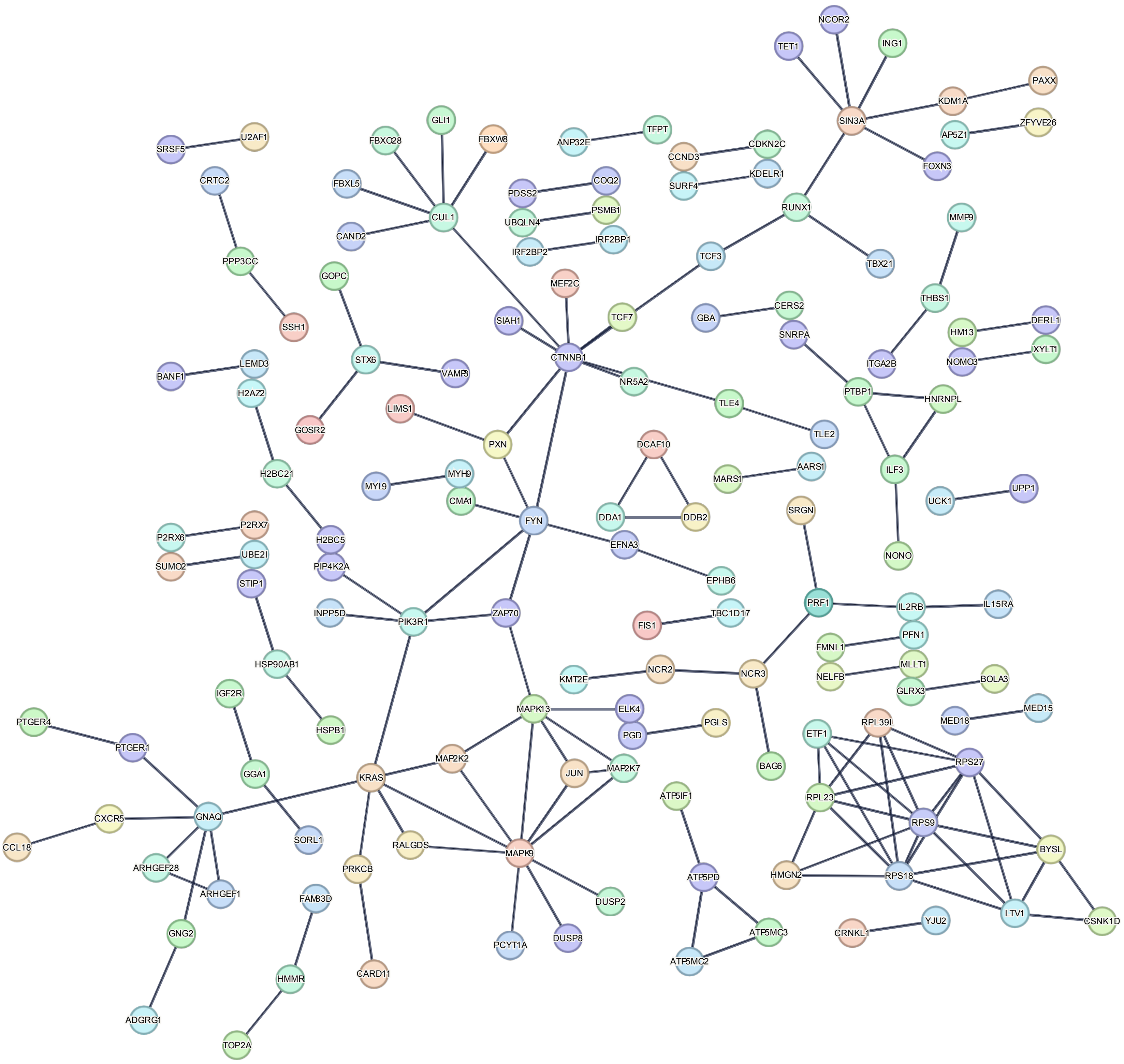
